# Empathy and the Structural Representation of Facial Affect: Evidence from a Genetic-Algorithm Face Synthesis Task

**DOI:** 10.64898/2026.06.02.729700

**Authors:** Bliss H. Cui, Peter J. Bex

## Abstract

Empathy has been linked to facial emotion recognition, but whether empathy is associated with the structural representation of facial affect (how observers position different affects relative to one another in face-shape space) remains largely unexplored. 53 adults completed a genetic-algorithm face task that generated prototypes for 13 affects using the Basel Face Model, and completed the 60-item Empathy Quotient (EQ). The genetic-algorithm task produced structurally distinct prototypes for all 13 affects (all paired tests p < .001 Bonferroni-corrected; Cohen’s dz 1.74–3.13), confirming that participants generated reliable, affect-specific face representations. A 13 × 13 between-affect distance matrix was then compared between higher-EQ (n = 30) and lower-EQ (n = 23) groups. 1 pair survived full correction across all 156 off-diagonal cells: Amusement × Contempt (Cohen’s d = −1.31), with higher-empathy participants representing these two affects as structurally closer to each other. Both amusement and contempt are social-evaluative affects that share overlapping facial action components, and this convergence may reflect heightened sensitivity to shared expressive structure among higher-empathy observers. In exploratory analyses, permutation testing and continuous-EQ correlations pointed to a broader pattern centered on social-evaluative affects (Amusement, Awe, Contempt, Fear, Happiness, Pride, Sadness, Interest). Individual differences in empathy appear most prominently associated with how social-evaluative affects are structurally positioned in face-shape space, suggesting that empathy modulates not just emotion recognition accuracy but the representational geometry of facial affect itself.

## 2. Introduction

Faces are a primary channel for the social communication of affect. Classic models of emotion recognition posit that observers assign facial expressions to a small number of discrete categories (Ekman, 1992), while dimensional models arrange affects along continuous axes such as valence and arousal (Russell, 1980). Behavioral and computational work over the past two decades has increasingly moved beyond both frameworks toward structural accounts in which affect categories occupy positions in a continuous representational space whose geometry may vary across observers (Barrett, Mesquita, & Gendron, 2011; Feldman Barrett, 2017; Barrett et al., 2019). On these accounts, the question of interest is not simply whether an observer can label an expression correctly, but how the observer’s internal representations of different affects are geometrically related to one another.

Parametric face models, combined with participant-driven search methods such as genetic algorithms, make it possible to characterize these internal representations with high precision. The Basel Face Model (Walker, Schönborn, Greifeneder, & Vetter, 2018; see also Walter & Bex, 2024) represents facial identity and expression as points in a compact 199-dimensional shape space derived from 3D face scans. When combined with a genetic-algorithm search procedure that explores this space efficiently (Carlisi et al., 2021; Binetti et al., 2022), the Basel Face Model can yield an observer’s internal prototype for a given affect: a single point in shape space that summarizes which facial configuration the observer treats as most representative of that affect. This approach is conceptually related to reverse-correlation techniques for recovering mental representations of faces (Dotsch & Todorov, 2012) and to multidimensional face-space models more broadly (Valentine, 1991), but it operates within a parametrically defined shape space rather than a noise-overlay or exemplar framework.

Empathy, the capacity to recognize and share others’ affective states, modulates several aspects of facial affect processing, including recognition accuracy, speed of detection, neural response amplitude, and spontaneous facial mimicry (Baron-Cohen & Wheelwright, 2004; Holland, O’Connell, & Dziobek, 2021; Adolphs, 2002). However, effect sizes linking self-reported empathy to categorical emotion recognition have been mixed: some studies report modest positive associations while others find effects close to zero (Olderbak & Wilhelm, 2017; Qiao et al., 2025). Most of this evidence comes from paradigms in which participants identify a small number of basic emotions from pre-selected stimulus sets. Whether empathy is also associated with the structural representation of affect, that is, with how different affect categories are positioned relative to one another in an observer’s representational geometry, has received little attention.

Recent work has demonstrated that genetic algorithms can recover observer-specific prototypes for emotional expressions and that substantial individual variability exists in these prototypes (Carlisi et al., 2021; Binetti et al., 2022). Follow-up studies have shown that the geometry of these prototypes (termed “expression perceptive fields”) predicts emotion categorization performance (Binetti et al., 2024), and that conceptual knowledge about emotions shapes the representational structure of facial emotion perception more broadly (Brooks & Freeman, 2018). However, no study to date has examined whether trait-level individual differences in empathy are associated with the structural geometry of facial affect representations in a parametric face-shape space.

The aim of the present study was to examine, in an exploratory framework, whether the Empathy Quotient is associated with the structural relations among participants’ generated facial affect prototypes. Participants completed a genetic-algorithm face task to synthesize 13 affects, a standardized set spanning both basic and social-evaluative categories (Amusement, Anger, Awe, Contempt, Disgust, Embarrassment, Fear, Happiness, Interest, Pride, Sadness, Shame, and Surprise), and completed the 60-item Empathy Quotient. Three analyses are reported: (1) task validation confirming that the 13 prototypes are structurally distinct; (2) a 13 × 13 between-affect distance matrix compared across empathy groups, with parametric correction across all 156 off-diagonal cells and a non-parametric permutation convergence check; and (3) continuous-EQ correlations across the same 156 pairs.

## 3. Methods

### 3.1 Participants

53 adults (40 women, 13 men; mean age 19.5 years, range 18–29) completed all study procedures and are included in the analyses reported here. Participants were recruited from the Northeastern University undergraduate research pool and were compensated with course credit. Inclusion required normal or corrected-to-normal vision by self report.

Participants additionally completed a depression questionnaire analyzed in a separate companion report; the two analyses share the same face-task data but use distinct dependent measures and address distinct questions.

The study was approved by the Northeastern University Institutional Review Board, and all participants provided written informed consent prior to taking part. This research was supported by the National Eye Institute under Award Number R01EY032162 (PI: Bex).

### 3.2 Apparatus and stimuli

Stimuli were displayed on a 27-inch iMac with a 5120 × 2880 (5K) resolution at a viewing distance of approximately 60 cm, subtending a visual angle of approximately 53° × 31°. The experiment was programmed and executed using MATLAB 2021a in conjunction with Psychtoolbox-3 (Brainard, 1997).

Face stimuli were parameterized using the 199-dimensional shape component of the Basel Face Model (Walker et al., 2018; see also Walter & Bex, 2024, for psychophysical applications of this model). Faces were rendered with the standard Basel viewpoint, lighting, and camera parameters, and presented at a fixed image scale on a uniform gray background.

### 3.3 Procedure: Genetic-algorithm face task

Each participant completed the genetic-algorithm face task for 13 affects presented in a randomized order: Amusement, Anger, Awe, Contempt, Disgust, Embarrassment, Fear, Happiness, Interest, Pride, Sadness, Shame, and Surprise. These 13 affects comprise a standardized set that spans both basic and social-evaluative categories of facial expression (Tracy & Robins, 2004). The genetic-algorithm procedure follows the approach described in Carlisi et al. (2021) and Binetti et al. (2022); additional psychophysical details of the Basel Face Model implementation used here are reported in Walter and Bex (2024). Each face image subtended approximately 6° of visual angle (screen width 60 cm at a viewing distance of 60 cm). For each affect the task ran for 7 generations of 12 faces per generation. Generations 1–6 were search generations; generation 7 was a final-selection round in which the participant chose a single face that best represented the affect (the “super face”). As described in §3.5, the analysis pipeline uses the median centroid across all selected faces rather than this single super face.

On each generation, participants viewed 12 new faces and selected between 0 and 12 of those faces that best represented the target affect; no minimum was imposed, ensuring that the search was entirely participant-led and free from experimenter bias. The composition of the next generation followed directly from the number of faces selected, s. If s = 0, the next generation consisted of 12 entirely new faces sampled from the Basel Face Model. If s = 1, 6 of the 12 faces were each produced by crossing the single parent face with a novel face (50% of the parent’s shape coefficients at random and 50% novel coefficients, with the specific half drawn independently per dimension), and the remaining 6 faces were novel. If s = 2, 6 of the 12 faces were produced by pairwise crossover of the 2 selected parents (50% from each), and the remaining 6 were novel. If s ≥ 3, 6 of the 12 offspring were produced by pairwise crossover between 2 parents drawn uniformly at random from the s selected faces (50% from each parent at random), and the remaining 6 were novel. Within-generation variability was maintained by the 6 novel faces included in every non-empty generation (see Figure 1 for a schematic). Trial structure, instructions, and timing were identical across affects.

**Figure 1.**
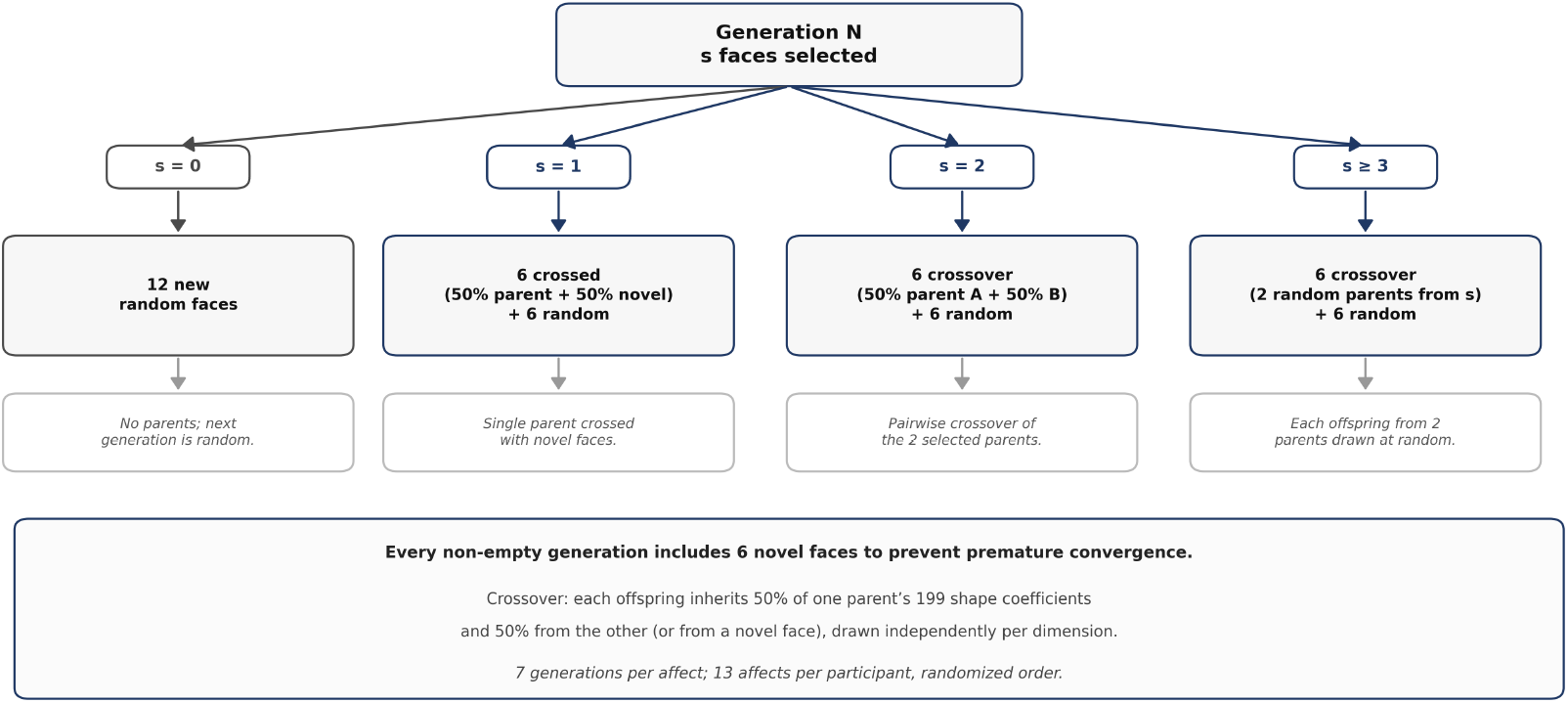
Schematic of the genetic-algorithm face task. Each generation presents 12 faces; the participant selects those matching the target affect. Selected faces are recombined to seed the next generation, while novel faces are introduced every generation to maintain variability. The process runs for 7 generations per affect.

### 3.4 Empathy measurement

Empathy was measured using the 60-item Empathy Quotient (EQ; Baron-Cohen & Wheelwright, 2004), which comprises 21 positively-keyed empathy items, 19 negatively-keyed empathy items, and 20 filler items. Positively-keyed items contributed 2 points for a “strongly agree” response and 1 point for “slightly agree” (0 for all other responses); negatively-keyed items contributed 2 points for “strongly disagree” and 1 point for “slightly disagree” (0 for all other responses); filler items were discarded. Total scores range from 0 to 80.

In this sample, EQ scores were approximately symmetric (M = 47.40, SD = 11.68, range 22–67; Shapiro–Wilk W indicated no departure from normality, p = .33, meaning the distribution of empathy scores was well-approximated by a normal curve). Empathy groups were defined via 1-dimensional k-means clustering with K = 2 applied directly to the 53 observed EQ scores. Centroids were initialized at the minimum and maximum observed scores, and the standard assignment–update cycle (reassign each score to the nearest centroid, then recompute each centroid as the mean of its assigned scores) was iterated to convergence. The data-driven split yielded a higher-empathy group (n = 30; M = 56.17, SD = 6.02, range 48–67) and a lower-empathy group (n = 23; M = 35.96, SD = 5.72, range 22–45). The empirical cluster boundary fell at EQ ≈ 46.1. The two groups differed on EQ by Welch’s t(48.6) = 12.46, Cohen’s d = 3.43. This data-driven procedure allows the group boundary to reflect the distribution of scores within this specific sample rather than relying on a fixed threshold (see Figure 4).

### 3.5 Analysis pipeline

#### 3.5.1 Per-participant, per-affect prototype (centroid)

For each participant and each of the 13 affects, we computed an affect-specific prototype defined as the element-wise median, across the 199 Basel Face Model shape dimensions, of all faces the participant selected across generations 1–7 for that affect. Each participant therefore contributed one 199-dimensional centroid per affect, yielding 13 centroids per participant. The number of selected faces contributing to a given centroid ranged from 2 to 44 across participants and affects (median = 16) from the 84 total faces each participant viewed for each affect; no participant–affect combination contributed fewer than 2 selected faces.

#### 3.5.2 Distance metric

Structural similarity between two 199-dimensional shape vectors A and B was quantified as the cosine distance D(A, B) = 1 − (A · B) / (‖A‖ × ‖B‖), which ranges from 0 (identical orientation) to 2 (exactly opposed orientation). Cosine distance is scale-invariant: it captures the orientation of a shape prototype within the 199-dimensional Basel space rather than its norm, making it appropriate for comparing Basel Face Model coefficient vectors across dimensions whose overall coefficient magnitudes may differ.

#### 3.5.3 Task validation

For each of the 13 affects, we computed a group-level prototype as the element-wise mean (arithmetic average) of the 53 per-participant centroids for that affect. For each participant– affect pair we then computed two distances: (a) a within-affect distance, defined as the cosine distance between the participant’s centroid for that affect and the group-level prototype of the same affect, and (b) a between-affect distance, defined as the mean cosine distance between the participant’s centroid for that affect and the group-level prototypes of the other 12 affects. A paired test of (between − within) was computed for each affect, together with Cohen’s dz and its 95% confidence interval (Figure 2). Multiple-comparison correction across the 13 affect-level tests was applied with both the Benjamini–Hochberg false discovery rate (BH-FDR) and the Bonferroni family-wise procedure.

**Figure 2.**
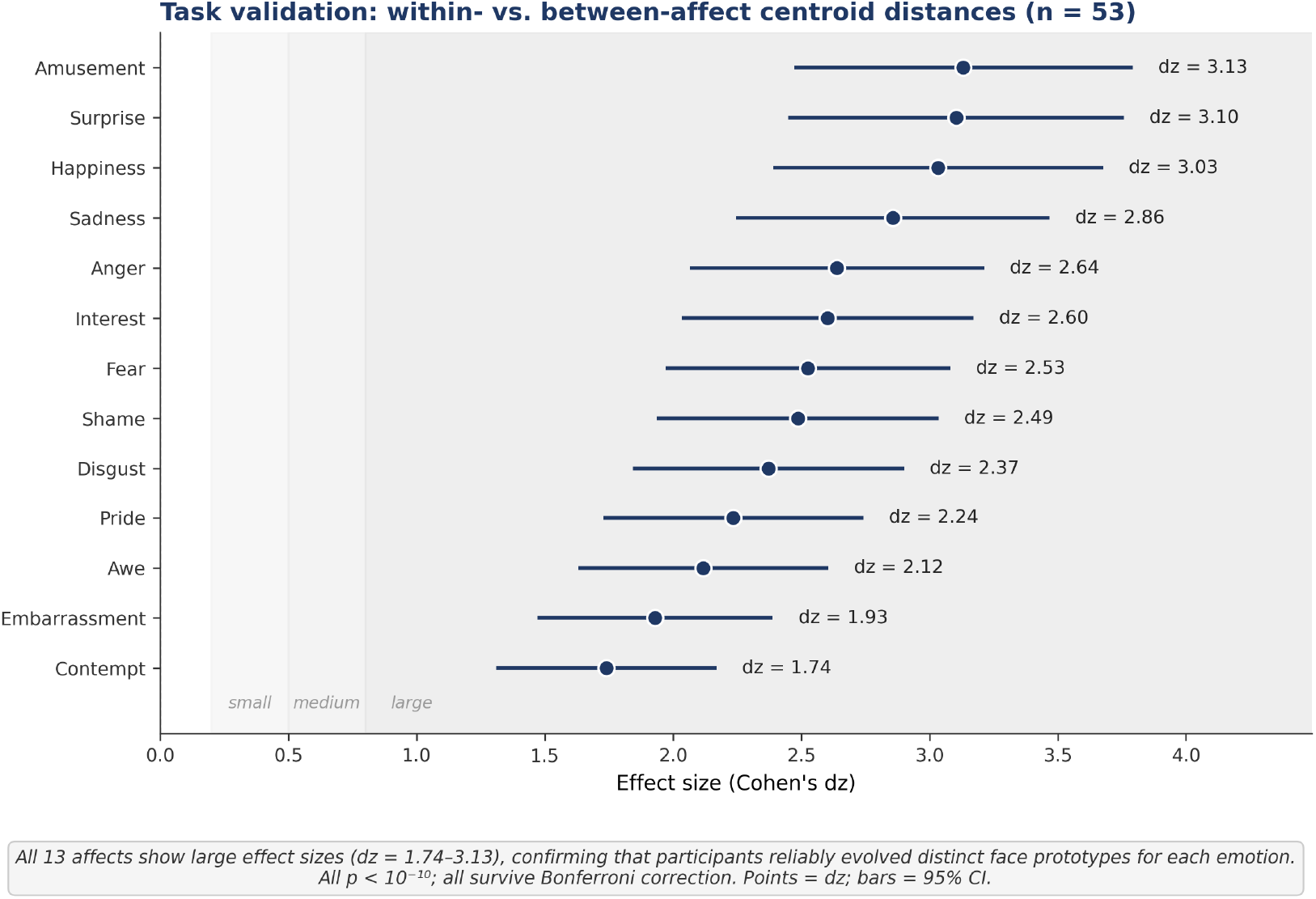
Task validation results. Cohen’s dz (with 95% CI) for the paired within-versus-between cosine distance test for each of the 13 affects. All 13 effects are significant after Bonferroni correction (p < .001). Dashed line marks dz = 0.8 (large effect).

#### 3.5.4 Empathy-group comparison: 13 × 13 between-affect distance matrix

For each ordered pair of affects (row affect e1, column affect e2) we computed, for every participant, the cosine distance from that participant’s e1 centroid to the mean e2 centroid of their own empathy group (i.e., the mean of the 199-dimensional e2 centroids across the other participants in the same group; leaving the target participant out). Welch’s two-sample t-test then compared these distances between the higher-empathy (n = 30) and lower-empathy (n = 23) groups at each of the 156 off-diagonal cells of the 13 × 13 matrix. Welch–Satterthwaite degrees of freedom are reported alongside each t. Multiple-comparison correction across the full 156-cell off-diagonal matrix was applied with both the Benjamini–Hochberg FDR and the Bonferroni family-wise procedure. The single cell that survives both procedures is reported as the headline empathy finding (Figure 3; Table 2). Effect sizes are reported as Cohen’s d with pooled standard deviation.

**Figure 3.**
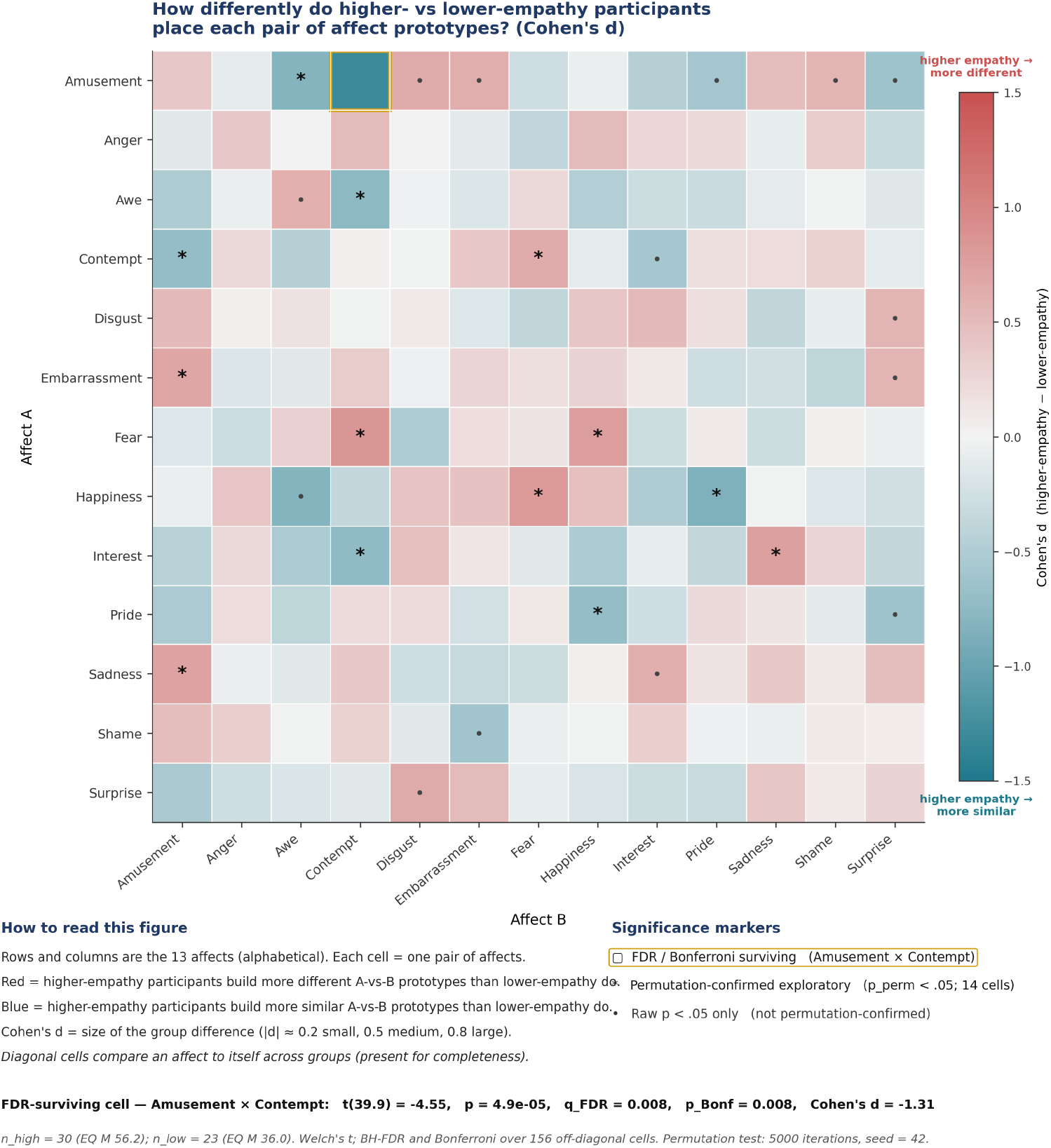
Empathy-group comparison across the 13 × 13 between-affect distance matrix. Each cell shows the signed mean difference (higher-EQ minus lower-EQ) in cosine distance. The Amusement × Contempt cell (starred) is the only pair surviving both BH-FDR and Bonferroni correction across all 156 off-diagonal cells.

**Figure 4.**
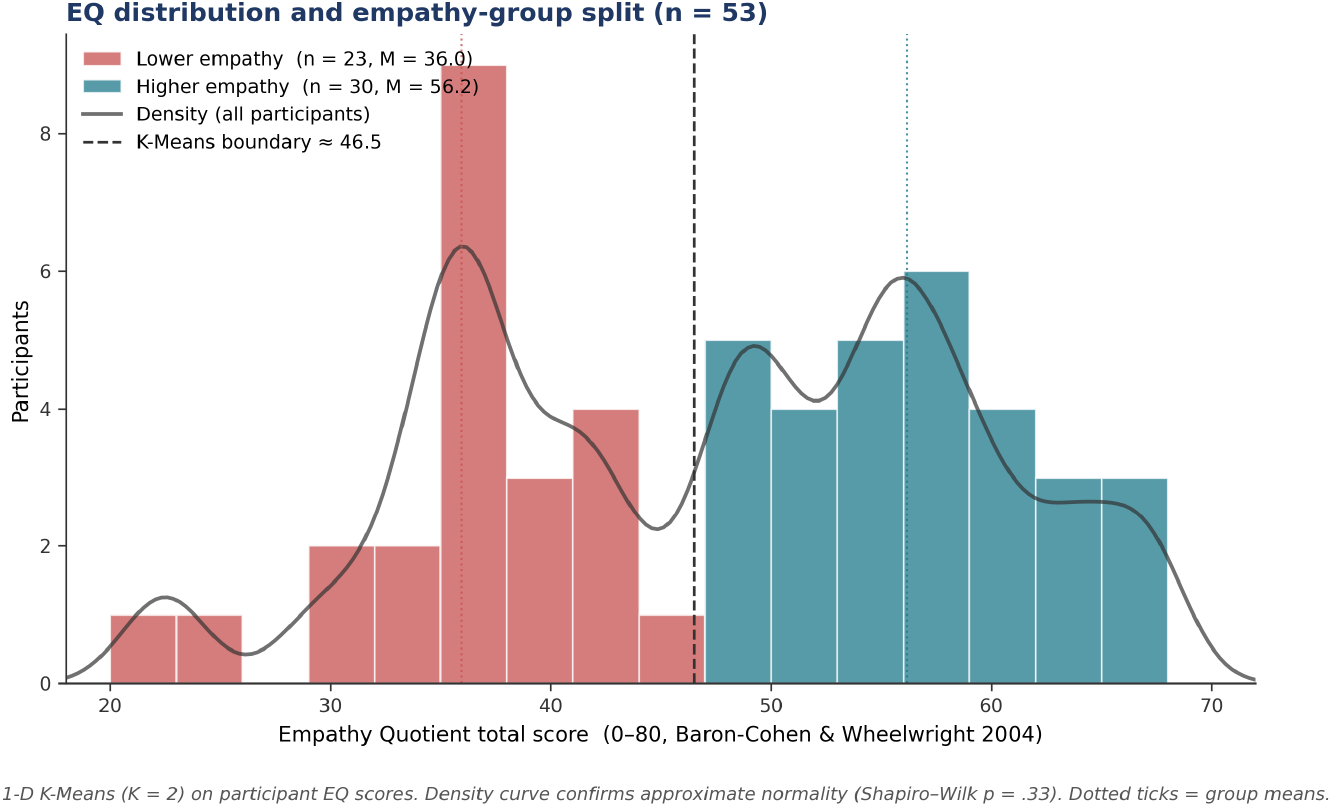
Distribution of Empathy Quotient (EQ) scores across the 53 participants. The dashed vertical line marks the empirical cluster boundary (EQ ≈ 46.1) from the 1-dimensional k-means split. Higher-empathy group: n = 30 (right); lower-empathy group: n = 23 (left).

#### 3.5.5 Permutation-based convergence check

As a non-parametric convergence check on the parametric matrix results, we ran a 5,000-iteration label-shuffle permutation test on each of the 28 cells for which the parametric test reached p < .05. Permutation testing was used because the Welch t-test assumes normally distributed group differences, and with unequal group sizes (30 vs. 23) across 156 cells, a non-parametric check ensures that the parametric p-values are not artifacts of distributional assumptions. On each iteration, empathy-group labels were shuffled without replacement across the 53 participants, the per-participant distances were regrouped under the shuffled labels, and the Welch t statistic at that cell was recomputed. The two-sided permutation p-value for a cell is the fraction of the 5,000 null iterations for which |t_null| ≥ |t_observed|. Cells for which p_perm < .05 are reported as exploratory secondary findings rather than as corrected-significant.

#### 3.5.6 Continuous-EQ correlations

To complement the group-based matrix analysis we also treated EQ as a continuous variable. For each of the 156 off-diagonal affect pairs (e1, e2) we computed a Pearson correlation across all 53 participants between each participant’s EQ total score and the cosine distance from their own e1 centroid to a leave-one-out group e2 prototype (the mean of the e2 centroids across the other 52 participants). Benjamini–Hochberg FDR correction was applied across the 156 correlations.

## 4. Results

### 4.1 Task validation: all 13 affect prototypes are structurally distinct

Each affect prototype generated by the genetic-algorithm task was structurally distinct from the group prototypes of the other 12 affects. The paired within-versus-between cosine distance test was significant for all 13 affects (all p < .001 after both Bonferroni and BH-FDR correction across the 13 tests). Cohen’s dz ranged from 1.74 (Contempt) to 3.13 (Amusement); 12 of the 13 affects exceeded dz = 1.9 (Table 1; Figure 2). These very large effect sizes indicate that participants produced reliable, affect-specific prototypes, establishing that the genetic-algorithm task is suitable for the empathy analyses that follow.

**Table 1.** Task validation: paired within-vs. between-affect cosine distance for each of the 13 affects (n = 53). Within M = mean cosine distance to the participant’s own-affect group prototype; Between M = mean cosine distance to the 12 other-affect group prototypes. All p-values survive both Bonferroni and BH-FDR correction.

| Affect | n | Within M | Between M | Diff | t | dz | p |
| --- | --- | --- | --- | --- | --- | --- | --- |
| Amusement | 53 | 0.800 | 1.000 | 0.199 | 22.80 | 3.13 | < .001 |
| Surprise | 53 | 0.815 | 0.994 | 0.179 | 22.59 | 3.10 | < .001 |
| Happiness | 53 | 0.782 | 0.997 | 0.214 | 22.09 | 3.03 | < .001 |
| Sadness | 53 | 0.808 | 1.010 | 0.202 | 20.80 | 2.86 | < .001 |
| Anger | 53 | 0.798 | 1.010 | 0.213 | 19.21 | 2.64 | < .001 |
| Interest | 53 | 0.816 | 0.998 | 0.182 | 18.95 | 2.60 | < .001 |
| Fear | 53 | 0.812 | 1.004 | 0.192 | 18.39 | 2.53 | < .001 |
| Shame | 53 | 0.811 | 1.010 | 0.199 | 18.10 | 2.49 | < .001 |
| Disgust | 53 | 0.826 | 1.005 | 0.179 | 17.27 | 2.37 | < .001 |
| Pride | 53 | 0.810 | 1.004 | 0.194 | 16.27 | 2.24 | < .001 |
| Awe | 53 | 0.830 | 0.998 | 0.168 | 15.42 | 2.12 | < .001 |
| Embarrassment | 53 | 0.838 | 1.003 | 0.165 | 14.05 | 1.93 | < .001 |
| Contempt | 53 | 0.861 | 1.000 | 0.138 | 12.66 | 1.74 | < .001 |

**Table 2.** All 27 off-diagonal cells reaching raw p < .05 in the empathy-group comparison. t = Welch’s t; p (raw) = uncorrected parametric p; Cohen’s d = pooled-SD effect size (negative = higher-empathy group closer); p (perm) = two-sided permutation p (5,000 iterations); Perm = permutation-confirmed at p < .05. Only Amusement × Contempt survives both BH-FDR and Bonferroni correction across all 156 off-diagonal cells.

| Affect pair | t | p (raw) | Cohen's d | p (perm) | Perm |
| --- | --- | --- | --- | --- | --- |
| Amusement × Contempt | -4.55 | 4.93e-05 | -1.31 | 0.008 | ✓ |
| Fear × Contempt | 3.13 | 0.003 | +0.84 | 0.017 | ✓ |
| Happiness × Pride | -3.09 | 0.003 | -0.85 | 0.038 | ✓ |
| Happiness × Awe | -2.96 | 0.005 | -0.81 | 0.065 |  |
| Amusement × Awe | -2.95 | 0.005 | -0.82 | 0.031 | ✓ |
| Happiness × Fear | 2.95 | 0.005 | +0.82 | 0.026 | ✓ |
| Awe × Contempt | -2.77 | 0.008 | -0.75 | 0.026 | ✓ |
| Fear × Happiness | 2.76 | 0.008 | +0.77 | 0.016 | ✓ |
| Interest × Sadness | 2.73 | 0.009 | +0.76 | 0.010 | ✓ |
| Sadness × Amusement | 2.68 | 0.010 | +0.73 | 0.021 | ✓ |
| Interest × Contempt | -2.64 | 0.011 | -0.73 | 0.044 | ✓ |
| Pride × Happiness | -2.57 | 0.013 | -0.70 | 0.028 | ✓ |
| Embarrassment × Amusement | 2.53 | 0.015 | +0.69 | 0.041 | ✓ |
| Contempt × Amusement | -2.46 | 0.018 | -0.70 | 0.013 | ✓ |
| Surprise × Disgust | 2.42 | 0.019 | +0.66 | 0.051 |  |
| Contempt × Fear | 2.38 | 0.021 | +0.67 | 0.020 | ✓ |
| Amusement × Surprise | -2.30 | 0.025 | -0.63 | 0.055 |  |
| Sadness × Interest | 2.29 | 0.026 | +0.62 | 0.054 |  |
| Amusement × Disgust | 2.26 | 0.030 | +0.66 | 0.063 |  |
| Amusement × Embarrassment | 2.22 | 0.032 | +0.63 | 0.110 |  |
| Pride × Surprise | -2.20 | 0.033 | -0.62 | 0.100 |  |
| Embarrassment × Surprise | 2.14 | 0.037 | +0.57 | 0.062 |  |
| Shame × Embarrassment | -2.13 | 0.039 | -0.60 | 0.078 |  |
| Contempt × Interest | -2.10 | 0.041 | -0.57 | 0.058 |  |
| Amusement × Shame | 2.06 | 0.044 | +0.57 | 0.168 |  |
| Amusement × Pride | -2.05 | 0.046 | -0.58 | 0.068 |  |
| Disgust × Surprise | 2.04 | 0.048 | +0.57 | 0.079 |  |

### 4.2 Empathy-group comparison: primary analysis

We compared the 13 × 13 matrix of between-affect cosine distances between the higher-EQ (n = 30) and lower-EQ (n = 23) groups. Across the 156 off-diagonal cells, 27 reached raw p < .05. After correction for multiple comparisons across all 156 cells, 1 cell survived both BH-FDR and Bonferroni correction: Amusement × Contempt (Welch’s t(39.9) = −4.55, p = 4.9 × 10^−5^, q_FDR = .008, p_Bonferroni = .008, Cohen’s d = −1.31).

The higher-empathy group represented Amusement and Contempt as structurally closer to each other (mean cosine distance = 0.947) than the lower-empathy group did (mean cosine distance = 1.030). The signed mean difference was −0.083, indicating that higher empathy is associated with a tighter geometric coupling between the amusement and contempt face prototypes in shape space. We treat this cell as the corrected-significance exemplar of a broader pattern in which empathy modulates the structural relations among social-evaluative affects (Figure 3; Table 2).

### 4.3 Permutation convergence: exploratory secondary findings

As a non-parametric convergence check, 5,000-iteration label-shuffle permutation tests were run on all 27 raw-significant matrix cells. 14 of the 27 cells survived permutation at p_perm < .05. These 14 pairs did not survive multiple-comparison correction across the full 156-cell matrix and are reported as exploratory.

The 14 permutation-confirmed cells involved 9 of the 13 affects: Amusement, Awe, Contempt, Embarrassment, Fear, Happiness, Interest, Pride, and Sadness. Contempt appeared in 6 of the 14 pairs and Amusement in 5, with additional involvement of Fear (4 pairs), Happiness (4 pairs), and the remaining affects in 1–2 pairs each. The pattern centers on the social-evaluative portion of the affect set. The full set of raw-significant and permutation results is reported in Table 2.

### 4.4 Continuous-EQ correlations: exploratory secondary findings

When EQ was treated as a continuous variable, 16 of the 156 off-diagonal affect pairs were correlated with EQ total score at raw p < .05. None survived BH-FDR correction across the 156 tests, so this analysis is reported as exploratory.

The strongest correlations were Happiness × Sadness (r = +.41, p = .003), indicating that higher EQ was associated with a greater structural separation between the happiness and sadness prototypes; Contempt × Shame (r = +.38, p = .006); and Amusement × Fear (r = −.33, p = .017), indicating that higher EQ was associated with a smaller separation between amusement and fear prototypes. The continuous-EQ analysis is broadly consistent with the social-evaluative pattern observed in the group comparison but does not, by itself, single out the Amusement × Contempt pair.

## 5. Discussion

### 5.1 Summary

The present study examined whether individual differences in empathy are associated with the structural geometry of facial affect representations. Three principal findings emerged. First, the genetic-algorithm face task produced structurally distinct prototypes for all 13 affects: every within-versus-between cosine distance test was significant after both Bonferroni and BH-FDR correction, with uniformly large effect sizes (dz = 1.74–3.13). Second, when a 13 × 13 between-affect distance matrix was compared across higher-empathy and lower-empathy groups, 1 cell survived correction across all 156 off-diagonal comparisons: higher-empathy participants represented Amusement and Contempt as structurally closer to each other than lower-empathy participants did (Cohen’s d = −1.31). Third, an exploratory pattern picked up by both permutation testing and continuous-EQ correlations pointed to social-evaluative affects (Amusement, Contempt, Fear, Happiness, Awe, Pride, Sadness, Interest) as the region of the affective space where empathy makes the most difference.

Taken together, these results suggest that individual differences in empathy are most prominently associated with how social-evaluative facial affects are structurally positioned relative to one another in face-shape space.

### 5.2 The Amusement–Contempt result

Amusement and Contempt are both social-evaluative affects: each involves an appraisal of another person’s behavior or social standing. Amusement typically signals positive social engagement (shared humor, affiliation, benign incongruity) while contempt signals a negative social evaluation, often involving perceived incompetence or moral transgression. Despite their opposing valence, the two expressions share overlapping facial action components, most notably asymmetric and suppressed smile elements (Ekman, 1992). Higher-empathy participants may be more attuned to this shared expressive structure, resulting in prototypes for amusement and contempt that are geometrically closer in the 199-dimensional shape space.

The effect size at this cell is large (d = −1.31), and it is the only pair that survives both BH-FDR and Bonferroni correction across the full 156-cell off-diagonal matrix. We treat it as the corrected-significance exemplar of a broader empathy/affect pattern rather than as a finding uniquely specific to amusement and contempt. The permutation-confirmed cells surrounding this pair (§4.3) reinforce this interpretation: the pattern is not isolated to a single cell but centers on the social-evaluative portion of the affect set.

### 5.3 The broader social-evaluative pattern

The 14 permutation-confirmed cells and the strongest continuous-EQ correlations both center on affects with a clear social-evaluative component: Amusement, Awe, Contempt, Embarrassment, Fear, Happiness, Interest, Pride, and Sadness. The four affects absent from this set (Anger, Disgust, Shame, and Surprise) are either more reflexive in nature (Surprise, Disgust; see Jack, Garrod, & Schyns, 2014) or less dependent on the appraisal of a specific social target (Anger, Shame). This is consistent with the broader literature framing empathy as a sensitivity to socially mediated affect rather than a uniform boost in emotional recognition across all categories (Baron-Cohen & Wheelwright, 2004; Adolphs, 2002), and aligns with evidence that the self-conscious emotions constitute a functionally distinct class tied to social appraisal (Tracy & Robins, 2004).

It is important to emphasize that the 14-cell exploratory pattern did not survive correction across the full 156-cell matrix. We report it because it converges with the headline finding and with the continuous-EQ analysis, and because the clustering of these cells around social-evaluative affects is consistent with theory. Confirmation in a larger, independently powered sample is needed before this broader pattern can be treated as established.

### 5.4 Relation to prior work

Most studies linking empathy to emotion perception have used forced-choice recognition of a small set of basic emotions. Meta-analytic evidence indicates that the association between self-reported empathy and categorical recognition accuracy is significant but modest (Olderbak & Wilhelm, 2017; Qiao et al., 2025), and may depend on which component of empathy is measured: empathic concern and personal distress, for example, appear to have opposing relationships with recognition performance (Israelashvili, Sauter, & Fischer, 2020). The present results suggest a complementary approach: empathy effects may be visible not only in categorical accuracy but also in the structural geometry of affect prototypes, that is, how an observer positions different affects relative to one another in a high-dimensional shape space. This structural perspective complements recognition-accuracy findings by revealing relationships between affect categories that categorical measures cannot capture.

The genetic-algorithm methodology used here builds on the work of Carlisi et al. (2021), who demonstrated that genetic algorithms combined with parametric face models can recover observer-specific emotion prototypes, and Binetti et al. (2022), who showed that individual differences in these prototypes predict recognition accuracy. Recent extensions have formalized the concept of “expression perceptive fields” (Binetti et al., 2024), which describe the tuning of an observer’s emotion representation within expression space. The present study complements this line of work by showing that trait empathy modulates the structural relations among these representations, not just their precision or category boundaries.

By including 7 self-conscious and social-evaluative affects alongside the 6 basic emotions, the present study also provides a broader view of the affective space than the canonical set of basic emotions alone (Ekman, 1992). The finding that empathy effects concentrate in the social-evaluative portion of this space is consistent with theoretical accounts that emphasize empathy’s role in interpersonal and social-evaluative processing (Baron-Cohen & Wheelwright, 2004; Holland et al., 2021) and aligns with evidence that conceptual knowledge shapes the representational structure of facial emotion perception (Brooks & Freeman, 2018). Empathy may function as a form of conceptual expertise that selectively refines the structural organization of social-evaluative affects, paralleling the way conceptual similarity shapes perceptual similarity in Brooks and Freeman’s framework.

### 5.5 Limitations

Several limitations should be noted. The sample comprised 53 undergraduate participants with a pronounced gender skew (40 women, 13 men), limiting generalizability. Group sizes of 30 and 23 provide adequate power for the headline finding (d = −1.31) but limit sensitivity for smaller effects; BH-FDR correction across 156 cells leaves only 1 surviving pair, even when many cells show effect sizes in the medium range. Stimuli varied only in the shape component of the Basel Face Model; conclusions are therefore limited to shape-based representations of affect and may not generalize to richer face stimuli that include textural cues. The Empathy Quotient is a self-report instrument, and its correspondence with behavioral measures of empathy is imperfect (Olderbak & Wilhelm, 2017). Finally, the genetic-algorithm task produces a population of selected faces per participant; the median-centroid summary discards within-participant variability that may itself carry information about how flexibly an observer represents a given affect.

### 5.6 Future directions

Replication with a larger and more demographically diverse sample would allow independent confirmation of the Amusement × Contempt finding and provide sufficient power to evaluate the broader social-evaluative pattern after full multiple-comparison correction. Adding texture variation to the Basel Face Model would test whether empathy effects persist or strengthen when textural cues are available alongside shape. Testing whether the pattern generalizes to other individual-difference measures, such as autistic traits, alexithymia, or behavioral empathy tasks (Israelashvili et al., 2020), would clarify whether the structural-geometry effect is specific to self-reported empathy or reflects a broader dimension of socio-emotional sensitivity. Finally, applying the expression perceptive field framework (Binetti et al., 2024) to datasets that include empathy measures could reveal whether the present geometric effects correspond to systematic differences in perceptive-field tuning.

## 6. Acknowledgments

This research was supported by the National Eye Institute under Award Number R01EY032162 (PI: Bex). We thank the members of the Bex laboratory who contributed to data collection and the Northeastern University undergraduate research pool participants.

